# Stasimon contributes to the loss of sensory synapses and motor neuron death in a mouse model of spinal muscular atrophy

**DOI:** 10.1101/608513

**Authors:** Christian M. Simon, Meaghan Van Alstyne, Francesco Lotti, Elena Bianchetti, Sarah Tisdale, George Z. Mentis, Livio Pellizzoni

**Author notes:** Corresponding author: Livio Pellizzoni, Center for Motor Neuron Biology and Disease, Department of Pathology and Cell Biology, Columbia University, 630 West 168^TH^ Street, New York, NY, 10032. Equal contribution (Christian M. Simon, Meaghan Van Alstyne).

## Abstract

Reduced expression of the SMN protein causes spinal muscular atrophy (SMA) – an inherited neurodegenerative disease characterized by multiple synaptic deficits and motor neuron loss. Here, we show that AAV9-mediated delivery of Stasimon – a gene encoding an ER-resident transmembrane protein regulated by SMN – improves motor function in a mouse model of SMA through multiple mechanisms. In proprioceptive neurons of SMA mice, Stasimon overexpression prevents the loss of afferent synapses on motor neurons and enhances sensory-motor neurotransmission. In SMA motor neurons, Stasimon suppresses the neurodegenerative process by selectively reducing phosphorylation but not upregulation of the tumor suppressor p53, both of which are converging events required to trigger neuronal death. We further show that Stasimon deficiency synergizes with SMA-related mechanisms of p53 upregulation to induce phosphorylation of p53. These findings identify Stasimon dysfunction induced by SMN deficiency as an upstream driver of cellular pathways that lead to synaptic loss and motor neuron degeneration, revealing a dual contribution of Stasimon to motor circuit pathology in SMA.

## Introduction

Spinal muscular atrophy (SMA) is an autosomal recessive neuromuscular disorder characterized by the progressive loss of spinal motor neurons and skeletal muscle atrophy (Burghes and Beattie, 2009; Tisdale and Pellizzoni, 2015). SMA is a consequence of ubiquitous reduction in the levels of the survival motor neuron (SMN) protein due to homozygous deletion or mutation of the *SMN1* gene with retention of the hypomorphic *SMN2* gene (Lefebvre et al., 1995). SMN has a well-characterized role in the assembly of small nuclear ribonucleoproteins (snRNPs) of the splicing machinery (Meister et al., 2001; Pellizzoni et al., 2002) as well as U7 snRNP that functions in 3’ end processing of histone mRNAs (Pillai et al., 2003; Tisdale et al., 2013). SMN has also been implicated in other aspects of RNA regulation including mRNA transport (Donlin-Asp et al., 2017). Consistent with its central role in RNA processing (Donlin-Asp et al., 2016; Li et al., 2014), SMN deficiency has been shown to induce widespread splicing dysregulation and transcriptome alterations in a variety of *in vivo* models (Bäumer et al., 2009; Doktor et al., 2017; Jangi et al., 2017; Zhang et al., 2008, 2013). The identification of downstream RNA targets of SMN deficiency that directly contribute to SMA pathology is of critical importance for elucidating disease mechanisms and revealing SMN-independent therapeutic approaches. To date, however, this has proven to be challenging due to the diversity of RNA pathways controlled by SMN and the complexity of SMA pathology in mouse models that more closely resemble the most severe form of the human disease.

Motor neurons are the cell type most severely affected by SMN deficiency, and their degeneration is a hallmark of SMA pathology (Burghes and Beattie, 2009; Tisdale and Pellizzoni, 2015). Importantly, selective genetic restoration of SMN in motor neurons of SMA mice has demonstrated that neuronal death is a cell autonomous process (Fletcher et al., 2017; Gogliotti et al., 2012; Martinez et al., 2012; McGovern et al., 2015). We previously demonstrated that activation of the tumor suppressor p53 is a key driver of motor neuron degeneration in the SMNΔ7 mouse model of SMA (Simon et al., 2017). We also showed that selectivity is established through the convergence of distinct mechanisms of p53 regulation including stabilization and phosphorylation of its N-terminal transcription-activation domain (TAD) (Simon et al., 2017), which only occur in the pool of vulnerable SMA motor neurons destined to die. Recently, we demonstrated that p53 upregulation results from dysregulated alternative splicing of Mdm2 and Mdm4 – the two main inhibitors of p53’s stability and function (Toledo and Wahl, 2006; Vousden and Prives, 2009) – due to reduced snRNP levels in SMA motor neurons (Van Alstyne et al., 2018a), thereby directly linking neurodegeneration to specific splicing changes induced by SMN deficiency. However, the converging mechanism(s) responsible for selective phosphorylation of the TAD of p53 required for degeneration of SMA motor neurons is unknown.

Characterization of SMA pathogenesis has identified multiple synaptic deficits in the motor circuit beyond motor neuron death that include dysfunction and loss of neuromuscular junctions (NMJ) as well as central proprioceptive sensory synapses onto motor neurons (Shorrock et al., 2019; Van Alstyne and Pellizzoni, 2016), which likely exert compounding effects on neuromuscular function. Studies in mouse models indicated that NMJ denervation occurs as a consequence of intrinsic effects of SMN deficiency in SMA motor neurons rather than skeletal muscle (Fletcher et al., 2017; Gavrilina et al., 2008; Iyer et al., 2015; Martinez et al., 2012) and that the pathogenic processes underlying NMJ denervation and death of motor neurons might be distinct (Kim et al., 2017; Van Alstyne et al., 2018a). The dysfunction and loss of proprioceptive synapses on somata and dendrites of SMA motor neurons is an early pathogenic event in SMA mice (Mentis et al., 2011), which is caused by the effects of SMN deficiency within proprioceptive neurons (Fletcher et al., 2017). The resulting reduction of excitatory drive from proprioceptive synapses induces hyperexcitability and reduces the firing of motor neurons contributing to impaired muscle contraction in SMA mice (Fletcher et al., 2017). Moreover, the deafferentation and dysfunction of motor neurons are mechanistically uncoupled from motor neuron death (Fletcher et al., 2017; Simon et al., 2016). However, while multiple insults to the synaptic integrity of sensory-motor circuits originate from defects in distinct neurons that coalesce to cause SMA pathology, the underlying SMN-dependent mechanisms are poorly understood.

This work aimed to address the outstanding questions described above by investigating the potential contribution of Stasimon dysfunction to SMA pathology in a mouse model of the disease. We previously identified Stasimon (also known as Tmem41b) as an evolutionarily conserved, U12 intron-containing gene regulated by SMN that is required for proper synaptic transmission in the motor circuit of *Drosophila* larvae and motor axon outgrowth in zebrafish (Lotti et al., 2012). Importantly, Stasimon contributes to motor circuit deficits in *Drosophila* and zebrafish models of SMA as its overexpression suppresses specific neuronal phenotypes induced by SMN deficiency in these model organisms (Lotti et al., 2012). Furthermore, we and others have found that SMN deficiency disrupts U12 splicing (Boulisfane et al., 2011; Doktor et al., 2017; Jangi et al., 2017; Lotti et al., 2012), and both misprocessing and reduced expression of Stasimon mRNA was found in disease-relevant motor circuit neurons of SMA mice (Lotti et al., 2012). Recent studies have also shown that Stasimon is a transmembrane protein localized to the ER that is essential for mouse embryonic development (Van Alstyne et al., 2018b) and involved in autophagy (Moretti et al., 2018; Morita et al., 2018; Shoemaker et al., 2019). However, Stasimon’s normal requirement in the mammalian motor circuit and a direct involvement in pathogenesis of SMA mouse models have not been established.

Here, we investigated the effects of Stasimon restoration in the SMNΔ7 mouse model of SMA by means of adeno-associated virus serotype 9 (AAV9)-mediated gene delivery. We found that overexpression of Stasimon rescues the loss of proprioceptive synapses on motor neurons, improves synaptic transmission, and suppresses motor neuron degeneration. Mechanistically, we demonstrate that the loss of Stasimon is sufficient to induce p53 phosphorylation, a key event underlying the selective degeneration of SMA motor neurons (Simon et al., 2017). Together, these results establish Stasimon’s direct involvement in distinct pathogenic cascades induced by SMN deficiency in proprioceptive neurons and motor neurons, revealing a dual contribution by Stasimon to sensory-motor circuit dysfunction in a mouse model of SMA.

## Results

### AAV9-mediated Stasimon gene delivery improves motor function in SMA mice

We sought to investigate the effects of increased expression of human Stasimon (abbreviated as STAS) on the phenotype of SMNΔ7 SMA mice by intracerebroventricular (ICV) injection of self-complementary AAV9 vectors at P0, an established method to study SMA disease mechanisms *in vivo* (Passini et al., 2010; Simon et al., 2017; Van Alstyne et al., 2018a). RT-qPCR analysis confirmed that injection of AAV9-STAS as well as AAV9-GFP and AAV9-SMN, which were used as negative and positive controls, respectively, resulted in robust expression of the corresponding mRNAs in the spinal cord of SMA mice at P11 (Figure 1A). AAV9-STAS did not increase full-length SMN2 mRNA or SMN protein levels in the spinal cords of SMA mice (Figure 1A and 1B). To rule out any potential indirect effects of AAV9-STAS on SMN function, we also looked at representative downstream targets of SMN-regulated RNA pathways. AAV9-STAS did not correct aberrant U12 splicing of endogenous Stasimon pre-mRNA (Lotti et al., 2012) or other RNA processing events such as U7-dependent histone H1c 3’ end misprocessing (Tisdale et al., 2013), p53-dependent induction of Cdkn1a mRNA (Simon et al., 2017) and reduced Chodl mRNA expression (Bäumer et al., 2009; Zhang et al., 2008) in the spinal cord of SMA mice at P11 (Figure 1C). In contrast, all of these SMN-dependent events were robustly corrected by AAV9-SMN as expected (Figure 1C). Consistent with Stasimon being a downstream mRNA target of U12 splicing dysfunction induced by SMN deficiency (Lotti et al., 2012), these results demonstrate that AAV9-STAS does not enhance SMN expression or function.

**Figure 1.**
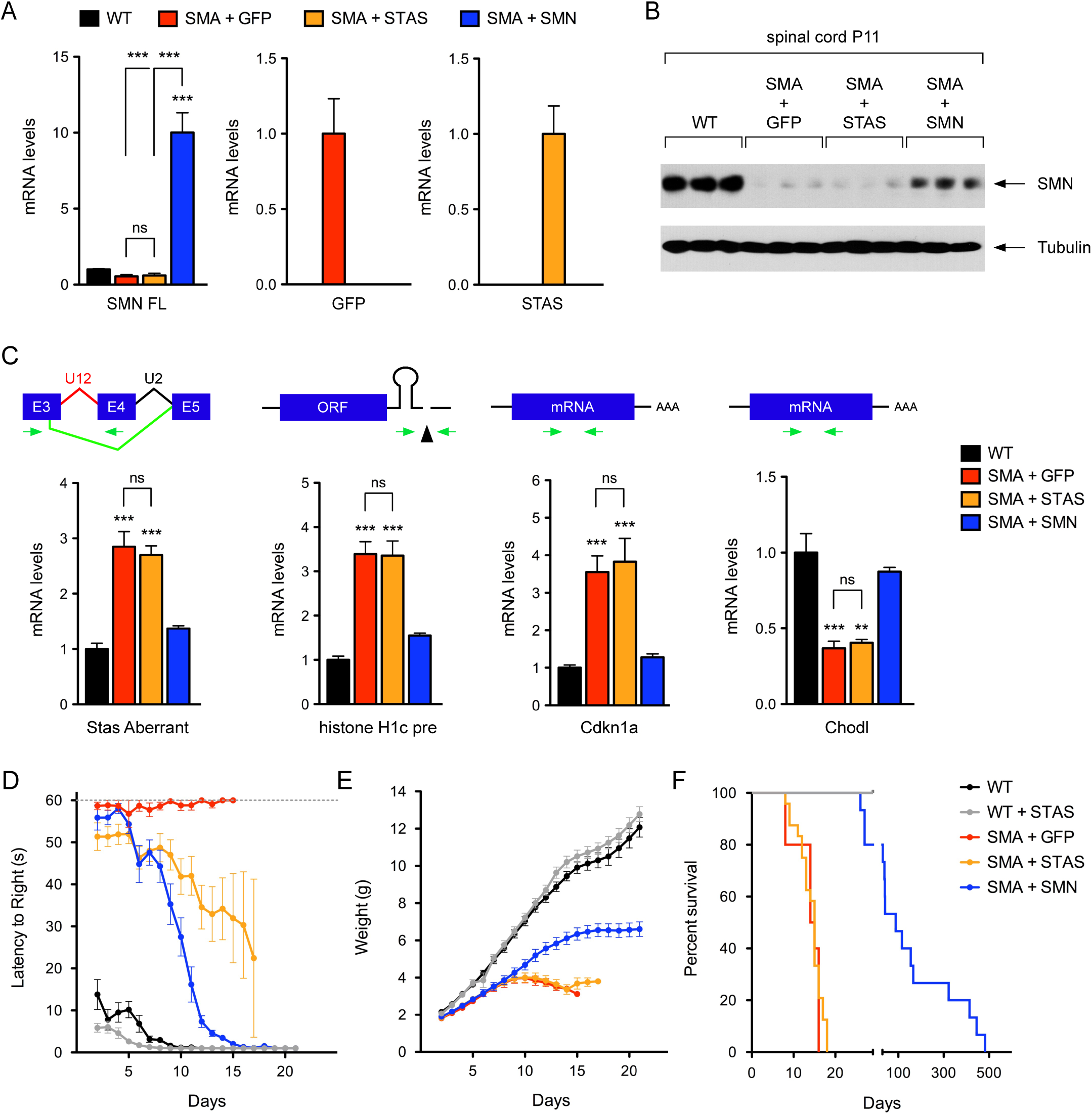
AAV9-mediated Stasimon gene delivery improves motor function in SMA mice. (A) RT-qPCR analysis of full-length human SMN (SMN FL), GFP, and human STAS mRNAs in the spinal cord of SMA mice injected with AAV9-GFP, AAV9-STAS or AAV9-SMN as well as uninjected WT mice at P11. Data represent mean and SEM (n=6). Statistics were performed with one-way ANOVA with Tukey’s *post hoc* test. *** P < 0.001; ns = not significant. (B) Western blot analysis of SMN protein levels in the spinal cord of AAV9-injected SMA mice and uninjected WT controls at P11. (C) RT-qPCR analysis of aberrantly spliced Stasimon mRNA, 3’ end-extended histone H1c mRNA, and total Cdkn1a and Chodl mRNAs in spinal cords from the same treatment groups as in (A) at P11. Schematics of the RNA processing events monitored in the assay are shown at the top. Data represent mean and SEM (n≥4). Statistics were performed with one-way ANOVA with Tukey’s *post hoc* test. ** P < 0.01; *** P < 0.001; ns = not significant. (D-F) Righting time (D), weight gain (E) and Kaplan-Meyer analysis of survival (F) of uninjected (n=10) and AAV9-STAS-injected (n=10) WT mice as well as SMA mice injected with AAV9-GFP (n=10), AAV9-STAS (n=24) and AAV9-SMN (n=15). Data represent mean and SEM.

We next investigated the effects of AAV9-STAS on the disease phenotype of SMA mice, which exhibit reduced weight, shortened life span and impaired motor behavior (Le et al., 2005). WT mice injected with AAV9-STAS showed no phenotypic abnormalities relative to uninjected WT littermates (Figure 1D-F), indicating no toxic effects of Stasimon overexpression *in vivo*. Furthermore, AAV9-SMN corrected all behavioral parameters and robustly extended survival in SMA as described previously (Foust et al., 2010; Passini et al., 2010). Remarkably, while AAV9-STAS had no effects on weight gain and survival, it improved motor behavior of SMA mice assessed by righting time (Figure 1D-F). Thus, Stasimon contributes to the motor phenotype of SMA mice, and the functional improvement elicited by AAV9-STAS occurs through SMN-independent mechanisms.

## AAV9-STAS rescues the loss of proprioceptive synapses on SMA motor neurons

To identify morphological improvements underlying the motor benefit of Stasimon restoration in SMA mice, we focused on synaptic connectivity of sensory-motor circuits controlling disease-relevant axial muscles. First, we analyzed the number of proprioceptive synapses onto motor neurons in the L1 lumbar spinal segment, which are strongly reduced in SMA mice (Mentis et al., 2011). To do so, we used immunohistochemistry with antibodies against the vesicular glutamate transporter 1 (VGluT1) to label proprioceptive synapses, and choline acetyltransferase (ChAT) as a motor neuron marker. SMA mice injected with AAV9-GFP exhibited a strong reduction of proprioceptive synapses onto somata and dendrites of L1 motor neurons at P11 relative to WT mice (Figure 2A, 2C and 2D). Strikingly, AAV9-STAS rescued the numbers of proprioceptive inputs on the somata and dendrites of motor neurons in SMA mice to levels similar to AAV9-SMN (Figure 2A, 2C and 2D).

**Figure 2.**
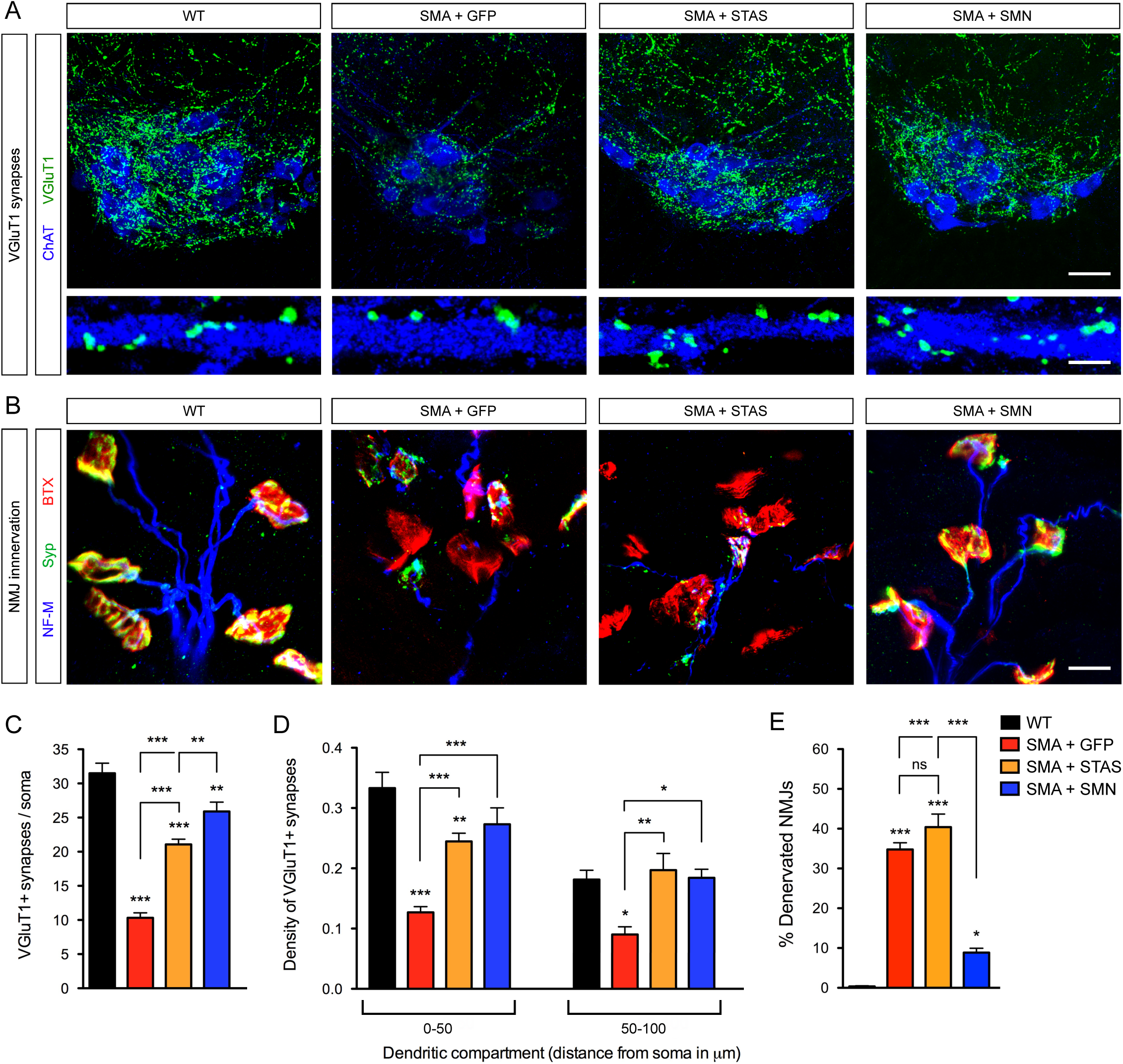
AAV9-STAS rescues the loss of proprioceptive synapses on SMA motor neurons. (A) Immunostaining of VGluT1+ synapses (green) and ChAT+ motor neurons (blue) in L1 spinal cord sections from SMA mice injected with AAV9-GFP, AAV9-STAS or AAV9-SMN as well as uninjected WT mice at P11. Scale bar = 50 µm. Lower panels show a magnified view of VGluT1 synaptic density on proximal dendrites. Scale bar = 10 µm. (B) NMJ staining with bungarotoxin (BTX, red), synaptophysin (Syp, green), and neurofilament (NF-M, blue) of QL muscles from the same groups as in (A) at P11. Scale bar = 15 µm. (C) Number of VGluT1+ synapses on the somata of L1 motor neurons from the same groups as in (A) at P11. Data represent mean and SEM from WT (n=22), SMA+AAV9-GFP (n=35), SMA+AAV9-STAS (n=32) and SMA+AAV9-SMN (n=21). Statistics were performed with one-way ANOVA with Tukey’s *post hoc* test. ** P < 0.01; *** P < 0.001. (D) Density of VGluT1+ synapses on the proximal dendrites of L1 motor neurons from the same groups as in (A) at P11. Two dendritic compartments were defined based on distance from the soma. Number of dendrites scored from the 0-50 µm compartment: WT (n=27), SMA+AAV9-GFP (n=21), SMA+AAV9-STAS (n=20) and SMA+AAV9-SMN (n=15). Number of dendrites scored from the 50-100 µm compartment: WT (n=12), SMA+AAV9-GFP (n=14), SMA+AAV9-STAS (n=10) and SMA+AAV9-SMN (n=8). Data represent mean and SEM. Statistics were performed with one-way ANOVA with Tukey’s *post hoc* test. * P < 0.05; ** P < 0.01; *** P < 0.001. (E) Percentage of fully denervated NMJs in the QL muscle from the same groups as in (A) at P11. Data represent mean and SEM (n=5). Statistics were performed with one-way ANOVA with Tukey’s *post hoc* test. * P < 0.05; *** P < 0.001; ns = not significant.

To examine the selectivity of Stasimon’s effects on synaptic loss, we investigated neuromuscular junction (NMJ) innervation of the quadratus lumborum (QL), a disease-relevant axial muscle innervated by vulnerable L1-L3 motor neurons that is severely denervated in SMA mice (Fletcher et al., 2017; Simon et al., 2017; Van Alstyne et al., 2018a). To do so, we used antibodies against neurofilament (NF-M) and synaptophysin (SYP) as presynaptic markers and fluorescently-labeled α-bungarotoxin (BTX) to label the acetylcholine receptor clusters on muscle fibers. At P11, NMJs of the QL in SMA animals injected with AAV9-GFP display ∼40% denervation in contrast to a fully innervated QL in WT mice (Figure 2B and 2E). Moreover, while AAV9-SMN injection restored NMJ innervation of the QL to near normal levels, AAV9-STAS did not have any beneficial effect on NMJ denervation in SMA mice (Figure 2B and 2E).

Taken together, these results reveal that Stasimon selectively restores the number of proprioceptive synapses in the sensory-motor circuit of SMA mice.

### Stasimon acts in proprioceptive neurons to improve sensory-motor synaptic transmission in SMA mice

We sought to determine whether restoration of proprioceptive synapses on SMA motor neurons by AAV9-STAS resulted in functional improvement of sensory-motor neurotransmission, which is severely disrupted in SMA mice (Fletcher et al., 2017; Mentis et al., 2011). To do so, we used the spinal cord *ex vivo* preparation and determined the amplitude of the sensory-motor monosynaptic reflex in the L1 spinal segment by simultaneous stimulation of the L1 dorsal root and recording from the ventral root of WT and SMA mice at P11 (Mentis et al., 2011). The amplitude of the monosynaptic reflex was reduced by ∼95% in SMA mice injected with AAV9-GFP and robustly enhanced upon SMN restoration with AAV9-SMN (Figure 3A and 3B). When we analyzed SMA mice injected with AAV9-STAS, we identified two distinct physiological responses in approximately equal proportions: a group of AAV9-STAS mice that exhibited a robust improvement of the monosynaptic reflex comparable to that of AAV9-SMN-injected SMA mice (Figure 3A and 3B), and another group that did not show increased response relative to AAV9-GFP treated SMA mice (Figure S1A). There were no significant differences between the two groups of SMA mice treated with AAV9-STAS in their body weight (Figure S1B), the levels of viral-mediated STAS mRNA overexpression in the spinal cord (Figure S1C), or the number of proprioceptive synapses on SMA motor neurons (aggregated data are shown in Figure 2) that could account for this observation. However, there was correlation with the motor performance of these SMA mice. We found that AAV9-STAS (R, righting) treated SMA mice that were able to right at the time of the assay (Figure S1D) had robust spinal reflex amplitudes (Figure 3A and 3B and Figure S1A), while AAV9-STAS (NR, non-righting) SMA mice with poor righting ability (Figure S1D) had small amplitudes (Figure S1A). Moreover, the levels of exon 7 containing, full-length SMN2 mRNA were higher in the spinal cord of AAV9-STAS (R) relative to AAV9-STAS (NR) SMA mice (Figure S1E). A marked reduction of SMN2 exon 7 splicing ensues at late symptomatic stages in SMA mice due to hypoxia and malnourishment, which exacerbate disease progression (Bebee et al., 2012; Sahashi et al., 2012). Thus, the observed differences in the electrophysiological responses of AAV9-STAS treated mice might reflect a degree of heterogeneity in the underlying severity of the SMA phenotype. Nevertheless, these experiments suggest that AAV9-STAS-mediated restoration of proprioceptive synapses can result in functional improvement of sensory-motor neurotransmission that correlates with enhanced motor function in SMA mice.

**Figure 3.**
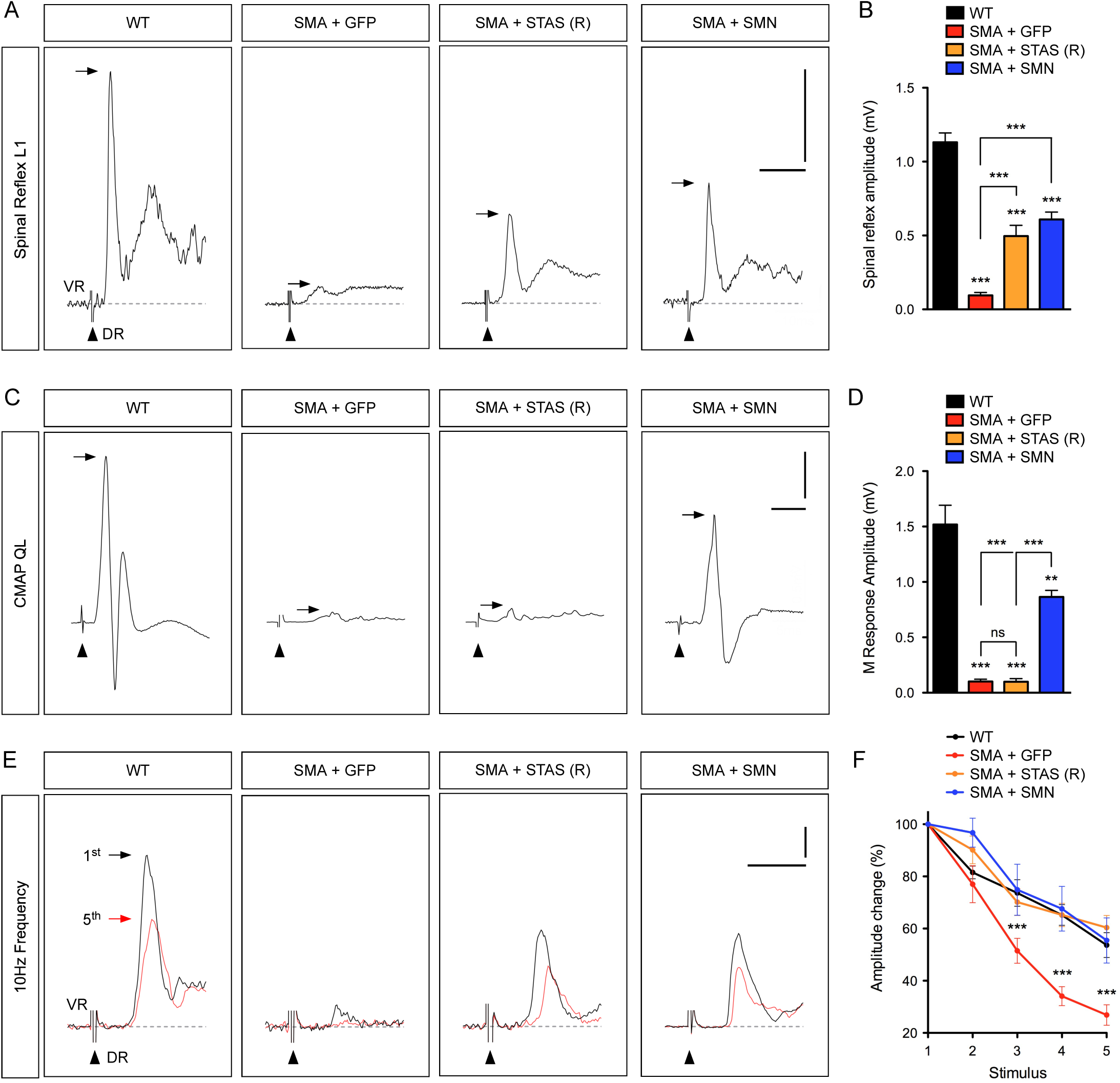
AAV9-STAS improves sensory-motor neurotransmission in SMA mice. (A) Representative traces of extracellular recordings from L1 ventral root (VR) following L1 dorsal root (DR) stimulation from SMA mice injected with AAV9-GFP, AAV9-STAS or AAV9-SMN as well as uninjected WT mice at P11. Arrows indicate the maximum amplitude of the monosynaptic reflex. Arrowheads indicate the stimulus artifact. Scale bars = 0.5 mV and 10 ms. (B) Quantification of spinal reflex amplitudes recorded from the same groups as in (A) at P11. Data represent mean and SEM (n≥4). Statistics were performed with one-way ANOVA with Tukey’s *post hoc* test. *** P < 0.001. (C) CMAP recordings from the QL muscle following L1 ventral root stimulation in the same groups as in (A) at P11. Arrows indicate maximum amplitude. Arrowheads indicate the stimulus artifact. Scale bars = 0.5 mV and 5 ms. (D) Quantification of M response amplitudes recorded from the QL muscle in the same groups as in (A) at P11. Data represent mean and SEM (n≥3). Statistics were performed with one-way ANOVA with Tukey’s *post hoc* test. ** P < 0.01; *** P < 0.001; ns = not significant. (E) Representative traces of the first (black) and fifth (red) VR responses recorded at P11 following stimulation of the homonymous L1 DR at 10 Hz from the same groups as in (A). Scale bars = 0.2 mV and 5 ms. (F) Quantification of amplitude changes in the monosynaptic reflex following 10 Hz stimulation from the same groups as in (A). Data represent mean and SEM (n≥3). Statistics were performed with two-way ANOVA with Tukey’s *post hoc* test. *** P < 0.001.

We next investigated NMJ function by stimulating motor axons in the L1 ventral root and recording the resultant compound muscle action potential (CMAP) in the QL muscle at P11. The CMAP of SMA mice injected with AAV9-GFP is strongly reduced compared to WT mice and significantly improved by AAV9-SMN injection (Figure 3C and 3D). AAV9-STAS-injected SMA mice did not show any improvement in the CMAP (Figure 3C and 3D), which was the case irrespective of their motor behavior or spinal reflex amplitudes at the time of the assay (Figure S1F), and consistent with the lack of effects on NMJ innervation (Figure 2B and 2E). Thus, the effects of Stasimon are selective for proprioceptive sensory synapses in SMA mice.

To dissect further the Stasimon-dependent functional improvement of the sensory motor circuit, we first determined the cellular contribution of motor neurons and proprioceptive synapses to the amplitude of the monosynaptic reflex in SMA mice. To do so, we restored SMN selectively in either proprioceptive neurons (SMA+SMN (PV-Cre)) or motor neurons (SMA+SMN (ChAT-Cre)) of SMA mice using a previously characterized conditional allele for Cre-dependent SMN restoration (Lutz et al., 2011). This approach rescues the number and function of proprioceptive neurons in SMA+SMN (PV-Cre) mice and motor neuron survival in SMA+SMN (ChAT-Cre), respectively (Fletcher et al., 2017). We found that motor neurons and proprioceptive neurons equally contribute to the reduction in the amplitude of the monosynaptic reflex of SMA mice (Figure S1G and S1H). To determine the reliability of neurotransmission of proprioceptive synapses irrespective of the number of motor neurons present, we performed repetitive stimulation of the sensory dorsal root L1 with simultaneous recording of the ventral root at high frequency (10 Hz) and compared the monosynaptic amplitude of the 1^st^ with the 5^th^ stimulus. We found a decrease of the monosynaptic amplitude of ∼40% in WT and ∼70% in SMA mice (Figure S1I and S1J), revealing higher synaptic depression in SMA mice likely due to impaired neurotransmission. While selective SMN restoration in motor neurons of SMA+SMN (ChAT-Cre) mice did not have any benefit, SMN expression in proprioceptive neurons of SMA+SMN (PV-Cre) mice reduced the synaptic depression of the monosynaptic reflex to WT levels, confirming that this experimental approach distinguishes the presynaptic contribution of proprioceptive neurotransmission in SMA (Figure S1I and S1J). Importantly, SMA mice injected with AAV9-STAS showed correction of synaptic depression to levels comparable to WT and AAV9-SMN treated SMA mice (Figure 3E and 3F). Collectively, these results indicate that AAV9-STAS ameliorates SMN-dependent sensory-motor circuit deficits by improving synaptic transmission in proprioceptive neurons of SMA mice.

### Stasimon contributes to motor neuron degeneration in SMA mice

Loss of motor neurons is a hallmark of SMA pathology (Burghes and Beattie, 2009; Tisdale and Pellizzoni, 2015) and we investigated whether Stasimon was implicated in this neurodegenerative process. To do so, we performed immunohistochemistry experiments and visualized motor neurons in the L1 and L5 spinal segments with antibodies against ChAT (Figure 4A and 4C). SMA mice injected with AAV-GFP displayed a loss of vulnerable L1 and L5 medial motor column (MMC) motor neurons at P11 (Figure 4B and 4D). Importantly, AAV9-STAS corrected this loss of SMA motor neurons with an efficacy similar to AAV9-SMN (Figure 4B and 4D). We recently showed that motor neuron death is caused by distinct, converging mechanisms of p53 activation in the SMNΔ7 mouse model (Simon et al., 2017; Van Alstyne et al., 2018a). To investigate whether Stasimon suppresses motor neuron death through the p53 pathway, we first immunostained against p53 and found no effects of AAV9-STAS on the number of p53+ L5 MMC motor neurons relative to AAV9-GFP treated SMA mice (Figure 5A and 5B), consistent with p53 upregulation being dependent on dysregulation of Mdm2 and Mdm4 alternative splicing (Van Alstyne et al., 2018a). Death of SMA neurons requires at least one other event converging on p53 that includes, but is not limited to, phosphorylation of p53 serine 18 (Simon et al., 2017) – a selective marker of degenerating SMA motor neurons. We found that AAV9-STAS significantly reduced the percentage of vulnerable L5 MMC motor neurons expressing phosphorylated p53^S18^ (p-p53^S18^) in SMA motor neurons (Figure 5C and 5D), consistent with Stasimon overexpression preventing SMA motor neuron death through modulation of p53 phosphorylation. Altogether, these experiments place Stasimon dysfunction upstream of the cascade of events driving phosphorylation of p53 and the subsequent process of motor neuron degeneration in SMA mice.

**Figure 4.**
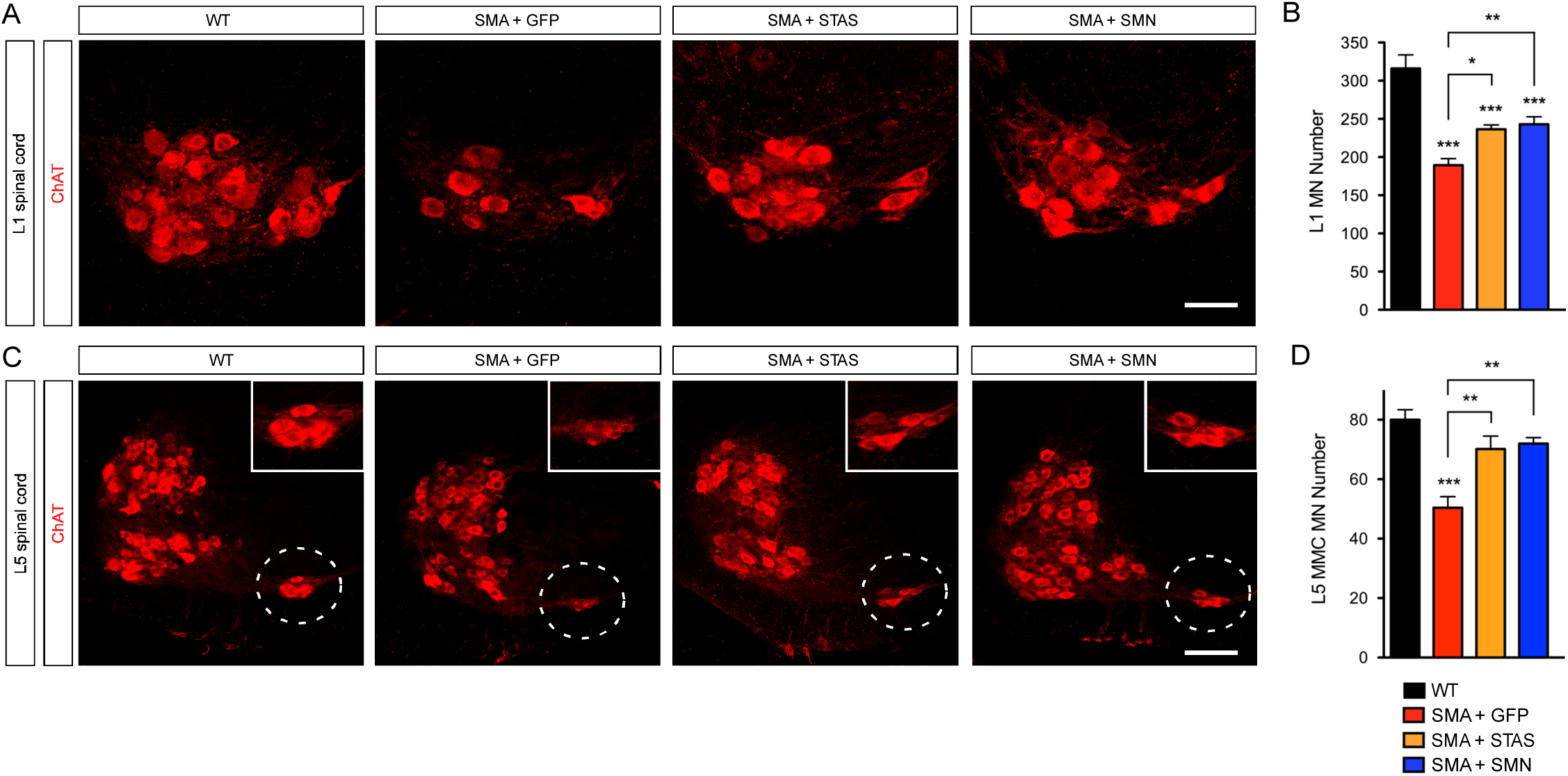
AAV9-STAS rescues vulnerable SMA motor neurons from degeneration. (A) ChAT immunostaining of L1 spinal cords from SMA mice injected with AAV9-GFP, AAV9-STAS or AAV9-SMN as well as uninjected WT mice at P11. Scale bar = 50 µm. (B) Total number of L1 motor neurons in the same groups as in (A) at P11. Data represent mean and SEM from WT (n=4), SMA+AAV9-GFP (n=6), SMA+AAV9-STAS (n=7) and SMA+AAV9-SMN (n=6). Statistics were performed with two-way ANOVA with Tukey’s *post hoc* test. * P < 0.05; ** P < 0.01; *** P < 0.001. (C) ChAT immunostaining of L5 spinal cords from the same groups as in (A) at P11. L5 MMC motor neurons are indicated by dotted circles and magnified in the insets. Scale bar = 125 µm. (D) Total number of L5 MMC motor neurons in the same groups as in (A) at P11. Data represent mean and SEM from WT (n=5), SMA+AAV9-GFP (n=5), SMA+AAV9-STAS (n=7) and SMA+AAV9-SMN (n=6). Statistics were performed with two-way ANOVA with Tukey’s *post hoc* test. ** P < 0.01; *** P < 0.001.

**Figure 5.**
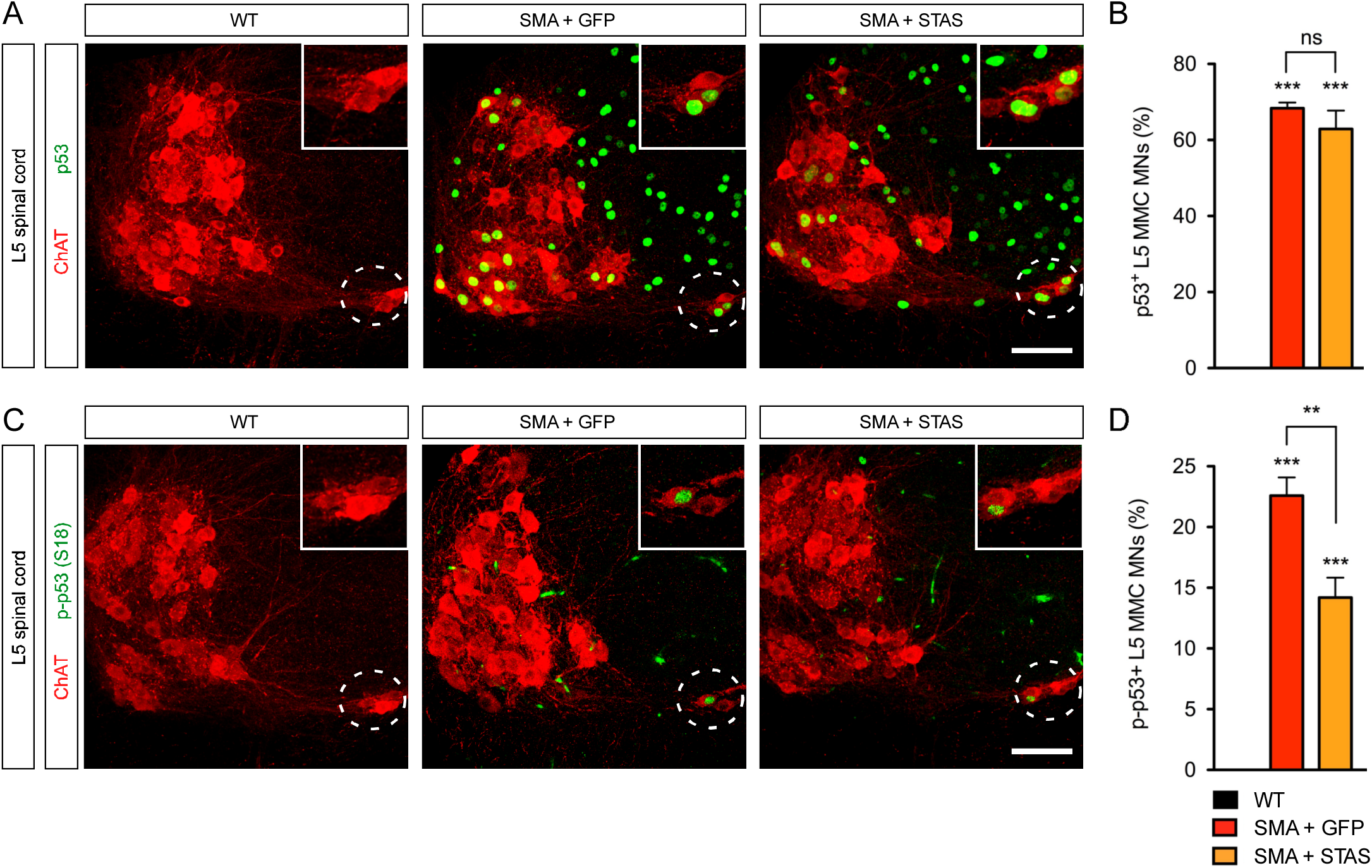
AAV9-STAS decreases p53^S18^ phosphorylation in vulnerable SMA motor neurons. (A) ChAT (red) and p53 (green) immunostaining of L5 spinal cords from uninjected WT mice and SMA mice injected with AAV9-GFP or AAV9-STAS at P11. L5 MMC motor neurons are indicated by dotted circles and magnified in the insets. Scale bar= 100 µm. (B) Percentage of p53+ L5 MMC motor neurons in the same groups as in (A). Data represent mean and SEM from WT (n=5), SMA+AAV9-GFP (n=5) and SMA+AAV9-STAS (n=5). Statistics were performed with one-way ANOVA with Tukey’s *post hoc* test. *** P < 0.001; ns = not significant. (C) ChAT (red) and p-p53^S18^ (green) immunostaining of L5 spinal cords from the same groups as in (A). L5 MMC motor neurons are indicated by dotted circles and magnified in the insets. Scale bar= 100 µm. (D) Percentage of p-p53^S18^+ L5 MMC motor neurons in the same groups as in (A). Data represent mean and SEM from WT (n=4), SMA+AAV9-GFP (n=5) and SMA+AAV9-STAS (n=6). Statistics were performed with one-way ANOVA with Tukey’s *post hoc* test. ** P < 0.01; *** P < 0.001.

## Stasimon deficiency is sufficient to induce p53S18 phosphorylation

Recent studies have shown that Stasimon is required for autophagy in cultured mammalian cells (Moretti et al., 2018; Morita et al., 2018; Shoemaker et al., 2019). Therefore, we sought to investigate the possible involvement of Stasimon-dependent autophagy dysregulation in SMA pathogenesis. First, we established a stable NIH3T3 cell line (NIH3T3-Stas_RNAi_) in which Stasimon mRNA levels are reduced to ∼10% of normal as monitored by RT-qPCR (Figure S2A). We then analyzed by Western blot the levels of the autophagy proteins p62 and LC3 in WT and Stas_RNAi_ NIH3T3 cells under basal conditions and following autophagy induction with serum starvation or inhibition by Bafilomycin A1 treatment (Figure S2B). We found that the levels of p62 and lipidated LC3 (LC3-II) were moderately increased in Stasimon-deficient relative to WT cells, consistent with dysregulation of autophagy (Moretti et al., 2018; Morita et al., 2018; Shoemaker et al., 2019). However, Western blot analysis showed no changes in the levels of both p62 and LC3 in spinal cords from WT and SMA mice at P11 (Figure S2C). Additionally, we performed immunohistochemistry experiments to investigate p62 expression in L1 motor neurons and proprioceptive neurons from L1 dorsal root ganglia (DRGs) of WT and SMA mice at P11 (Figure S2D and S2E), but did not find any major difference in the levels and localization of p62 induced by SMN deficiency in these disease-relevant neurons *in vivo*. These results suggest that autophagy dysregulation is unlikely to underlie the Stasimon-dependent deficits in SMA motor circuit.

Next, we sought to determine whether loss of Stasimon is sufficient to induce p53^S18^ phosphorylation as suggested by the reduction of p-p53^S18^ in SMA motor neurons described above. We have previously shown that upregulation but not N-terminal phosphorylation of p53 in SMA motor neurons is due to dysregulation of Mdm2 and Mdm4 alternative splicing (Van Alstyne et al., 2018a). Therefore, we devised a cell-based assay to investigate the role of Stasimon deficiency on p-p53^S18^ phosphorylation following p53 anti-repression induced by modulation of Mdm2 and Mdm4 splicing (Figure 6A). To do so, we treated WT and Stas_RNAi_ NIH3T3 cells either with control morpholino oligonucleotides (MOs) or with splice-switching MOs that induce selective skipping of Mdm2 exon 3 and Mdm4 exon 7 as previously reported (Figure 6B) (Van Alstyne et al., 2018a). We then performed immunofluorescence analysis and determined the percentage of treated NIH3T3 cells with nuclear accumulation of either p53 or p-p53^S18^. Treatment with Mdm2/4 MOs induced strong stabilization and nuclear accumulation of p53 in nearly all WT and Stas_RNAi_ NIH3T3 cells (Figure 6C and 6D), while there was no detectable nuclear p53 in cells treated with control MO. Importantly, we found that p53^S18^ was phosphorylated in nearly 80% of Stas_RNAi_NIH3T3 cells treated with Mdm2/4 MOs (Figure 6E and 6F), while little if any p-p53^S18^ was detected in all other treatment groups. These results show that Stasimon deficiency does not induce stabilization and nuclear accumulation of p53 in mammalian cells; however, loss of Stasimon is sufficient to induce N-terminal phosphorylation of p53 when converging with pre-existing mechanisms of p53 upregulation, mechanistically paralleling the paradigm driving degeneration of SMA motor neurons.

**Figure 6.**
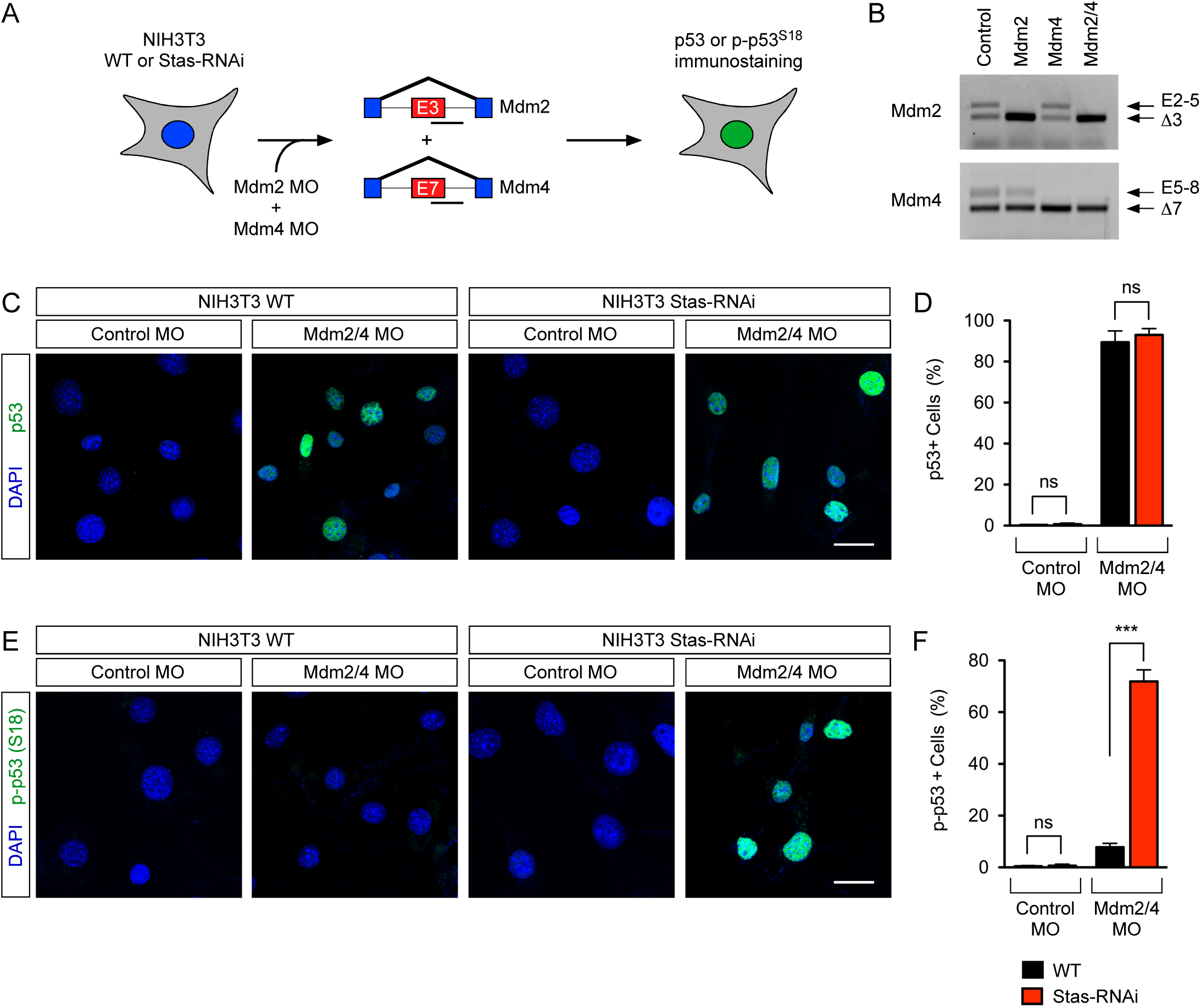
Stasimon deficiency is sufficient to induce p53^S18^ phosphorylation but not upregulation of p53 in mammalian cells. (A) Schematic of the experimental design. NIH3T3 cells (WT or Stas_RNAi_) were treated either with control MOs or with splice-switching antisense MOs targeting the 5’ splice sites of Mdm2 exon 3 and Mdm4 exon 7. Nuclear accumulation of p53 and p-p53^S18^ was then assessed by immunofluorescence analysis. (B) RT-PCR analysis of Mdm2 and Mdm4 alternative splicing in NIH3T3 cells treated with control MOs as well as Mdm2 and Mdm4 MOs either alone or in combination. (C) Immunofluorescence analysis of total p53 (green) in WT and Stas_RNAi_ NIH3T3 cells treated with the indicated MOs. Nuclei were counterstained with DAPI (blue). Scale bar= 25 µm. (D) Percentage of p53+ NIH3T3 cells from the same treatment groups as in (C). Data represent mean and SEM (n≥3). Statistics were performed with two-tailed unpaired Student’s t-test. ns = not significant. (E) Immunofluorescence analysis of p-p53^S18^ (green) in WT and Stas_RNAi_ NIH3T3 cells treated with the indicated MOs. Nuclei were counterstained with DAPI (blue). Scale bar= 25 µm. (F) Percentage of p-p53^S18^+ NIH3T3 cells from the same treatment groups as in (E). Data represent mean and SEM (n≥3). Statistics were performed with two-tailed unpaired Student’s t-test. *** P < 0.001; ns = not significant.

## Discussion

Motor dysfunction in SMA involves integration of multiple pathogenic insults induced by SMN deficiency in distinct neurons of the motor circuit (Shorrock et al., 2019; Van Alstyne and Pellizzoni, 2016). SMN has key roles in several RNA pathways that control gene expression (Donlin-Asp et al., 2016; Li et al., 2014) and establishing causal links between dysregulation of downstream mRNA targets of SMN function and SMA pathogenesis is fundamental for understanding disease mechanisms as well as identifying genes and pathways relevant for sensory-motor function that may also represent therapeutic targets. Here we show that AAV9-mediated Stasimon gene delivery improves motor function and corrects select pathogenic events in the motor circuit of SMA mice. Stasimon overexpression prevents the loss of proprioceptive synapses on somata and dendrites of SMA motor neurons with consequent functional improvement of sensory-motor neurotransmission that correlates with improved motor function. Stasimon restoration also rescues vulnerable motor neurons from degeneration, which we have previously shown to involve distinct mechanisms of p53 activation including upregulation and phosphorylation of the N-terminal TAD of p53 (Simon et al., 2017). Furthermore, our results indicate that Stasimon dysfunction induced by SMN deficiency is the upstream trigger of a signaling cascade(s) converging on p53 to drive its phosphorylation and the subsequent death of vulnerable motor neurons in SMA mice. Taken together, these findings reveal the direct contribution of Stasimon to distinct deficits in the sensory-motor circuit of SMA mice and highlight an unexpected link between Stasimon dysfunction and p53-dependent mechanisms of neurodegeneration. More broadly, they are consistent with a key role of Stasimon in development and function of the mammalian motor system.

Our findings that Stasimon restoration improves some but not all of the disease-related deficits in SMA mice supports the emerging view that a composite set of defects involving SMN-dependent RNA splicing alterations in specific genes and possibly disruption of other RNA pathways contribute to SMA pathology. We previously discovered Stasimon as a U12-intron containing gene regulated by SMN whose dysfunction results in specific neuronal deficits in the motor circuit of *Drosophila* and zebrafish models of SMA (Lotti et al., 2012). We determine here the contribution of Stasimon to SMA pathology in a mouse model that more accurately recapitulates genetic and phenotypic aspects of the human disease, establishing Stasimon as a disease-relevant target of SMN deficiency across evolution. Furthermore, as Stasimon is a downstream target of U12 splicing dysfunction induced by SMN deficiency (Lotti et al., 2012), this study strengthens the proposed pathogenic role of minor splicing disruption and the central contribution of Stasimon as an effector of U12-dependent motor circuit deficits in SMA, expanding our understanding of the RNA-mediated mechanisms of the disease.

Based on our findings and earlier genetic studies in SMA mice, it is plausible to draw some conclusions with regard to the spatial contribution of Stasimon to motor circuit pathology (Figure 7). On one hand, selective restoration of SMN in motor neurons indicated that induction of cell death is cell autonomous (Fletcher et al., 2017; Gogliotti et al., 2012; Martinez et al., 2012; McGovern et al., 2015), suggesting that the effects of AAV9-STAS on SMA motor neuron survival are mediated by Stasimon restoration in these cells. On the other hand, deafferentation is caused by SMN deficiency in proprioceptive neurons (Fletcher et al., 2017) and our results directly implicate Stasimon dysfunction in this process. Additionally, and in agreement with the spatial requirement of SMN and Stasimon in the *Drosophila* motor circuit (Imlach et al., 2012; Lotti et al., 2012), our studies of synaptic depression indicate that Stasimon acts in proprioceptive neurons to improve the function of sensory-motor synapses. Given that silencing proprioceptive neurotransmission alone does not induce synaptic loss (Fletcher et al., 2017), it is possible that synaptic dysfunction and deafferentation may result from disruption of distinct Stasimon-dependent events. These events are also different from those underlying the loss of NMJ synapses as indicated by the lack of improvement of neuromuscular innervation by Stasimon overexpression in SMA mice, which highlights diversity in the mechanisms of synaptic loss induced by SMN deficiency. Consistent with previous studies (Kim et al., 2017; Van Alstyne et al., 2018a), the protective effect of Stasimon on motor neuron survival but not NMJ innervation corroborates the view that these pathogenic hallmarks of SMA while both originating in motor neurons are mechanistically uncoupled.

**Figure 7.**
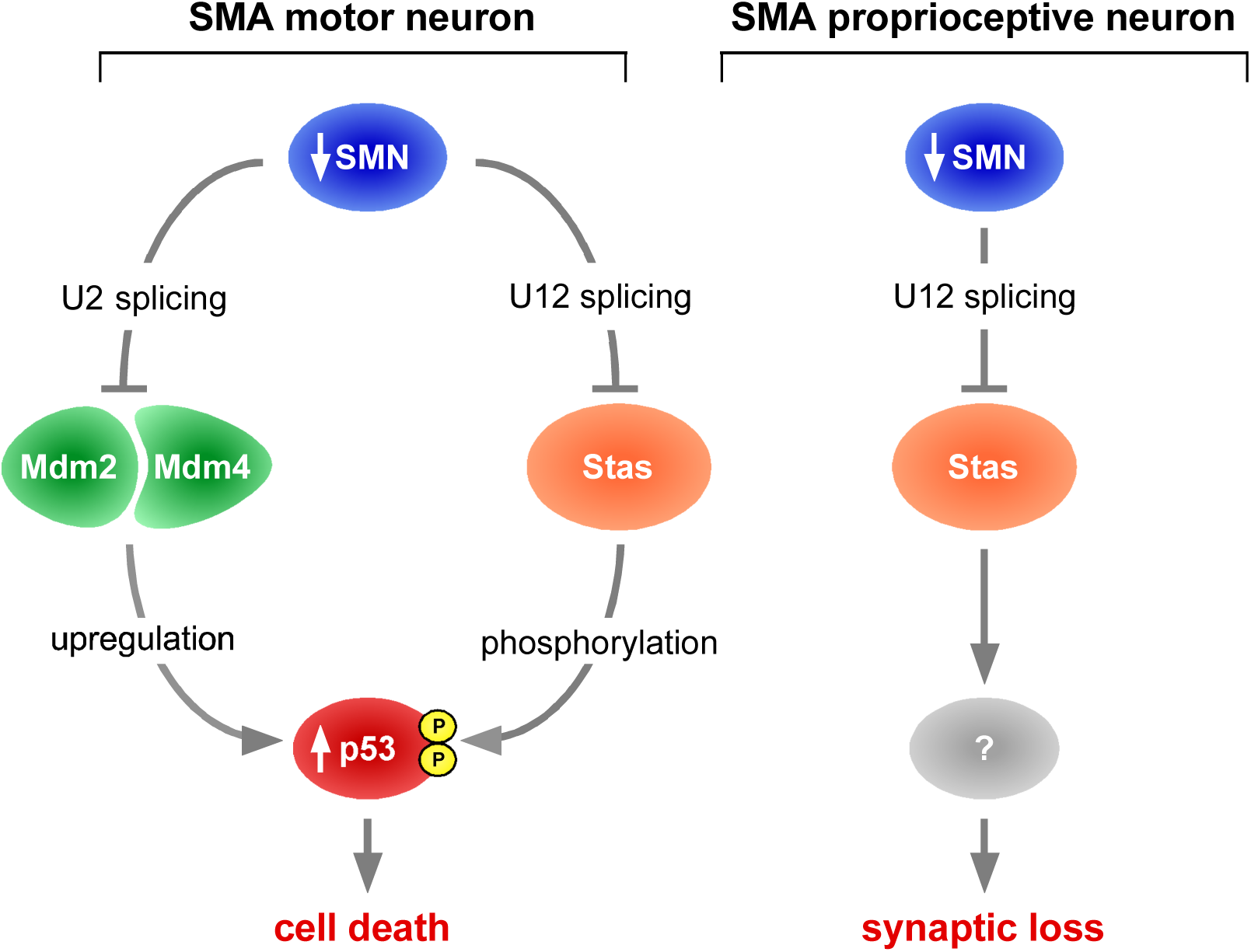
Dual contribution of Stasimon to sensory-motor circuit pathology in SMA. Schematic of the role of Stasimon in the degeneration of motor neurons and the loss of proprioceptive synapses in SMA (see text for further details).

Recent studies have shown that Stasimon is an ER-resident six-transmembrane pass protein that is essential for mouse embryonic development (Van Alstyne et al., 2018b) and functions in autophagy (Moretti et al., 2018; Morita et al., 2018; Shoemaker et al., 2019). However, we found no evidence for overt autophagy dysregulation in motor circuit neurons of SMA mice. Furthermore, Stasimon is enriched at ER-mitochondrial contact sites (Van Alstyne et al., 2018b), which are implicated in a variety of biological processes beyond autophagy that are important for neuronal physiology and disease (Paillusson et al., 2016). Elucidation of the downstream molecular mechanisms underlying dysfunction and loss of proprioceptive synapses in SMA will require greater knowledge of the full breadth of Stasimon functions. It will also need to integrate previous, seemingly unrelated findings implicating dysregulation of endocytosis (Ackermann et al., 2013; Hosseinibarkooie et al., 2016; Janzen et al., 2018; Riessland et al., 2017) and UBA1/GARS-dependent pathways (Shorrock et al., 2018) in the loss of SMA proprioceptive synapses into a coherent model.

The observation that Stasimon restoration suppresses motor neuron degeneration was unexpected, and we sought a mechanistic explanation in the context of the current understanding of the motor neuron death pathway. Based on previous studies in the SMNΔ7 mouse model of SMA (Simon et al., 2017), we have proposed a model in which selective degeneration of vulnerable SMA motor neurons requires the convergence of p53 stabilization with one or more additional events, which include post-translational modifications of p53 such as phosphorylation of its N-terminal TAD that specifically direct the p53 response towards degeneration (Figure 7). Recently, we demonstrated that dysregulation of Mdm2 and Mdm4 alternative splicing induced by SMN-dependent snRNP dysfunction underlies the upregulation of p53 in SMA motor neurons (Van Alstyne et al., 2018a). We also showed that these events are necessary but not sufficient to induce motor neuron death *in vivo* (Van Alstyne et al., 2018a), leaving open the question as to which SMN function(s) and downstream targets are involved in the other branch of the p53-dependent motor neuron death pathway. Our findings that AAV9-STAS reduces phosphorylation of p53^S18^ and rescues survival in SMA motor neurons *in vivo* together with the observation that Stasimon deficiency is sufficient to induce p53^S18^ phosphorylation *in vitro* fill this knowledge gap and place Stasimon dysfunction as an upstream driver of the neurodegenerative cascade. Since degeneration of SMA motor neurons requires phosphorylation of p53 at multiple residues beyond S18 (Simon et al., 2017), Stasimon likely triggers some of these additional p53 modifications required for neurodegeneration, but the lack of specific reagents prevents their identification at this time. Future studies will also be needed to determine whether Stasimon deficiency is sufficient to drive SMA motor neuron death when converging with Mdm2-and Mdm4-mediated p53 upregulation.

In summary, our work identifies Stasimon as a key determinant of defective synaptic connectivity and function in the SMA sensory-motor circuit as well as an upstream trigger of a signaling cascade(s) that feeds into p53 and the motor neuron cell death pathway. SMN deficiency has previously been linked to the activation of several kinases implicated in the degeneration of SMA motor neurons (Genabai et al., 2015; Kannan et al., 2018; Miller et al., 2015; Ng et al., 2015) and known to phosphorylate p53 (Toledo and Wahl, 2006; Vousden and Prives, 2009). Identification of the Stasimon-dependent signal transduction pathways and kinase(s) that drive p53 phosphorylation and motor neuron death in SMA might reveal viable targets for the development of neuroprotective agents for SMA therapy. Thus, further characterization of Stasimon biology will not only provide critical insight into the mechanisms underlying normal development and disease of the motor circuit but may also yield novel targets for developing SMN-independent SMA therapeutics.

## Material and Methods

### DNA cloning and viral production

For AAV9 production, DNA fragments corresponding to the open reading frames of GFP, human SMN (NM_000344), and human STAS (NM_015012) were generated by PCR using plasmid templates obtained from OriGene and cloned downstream of the CMV enhancer and chicken beta-actin (CB) hybrid promoter in the dsAAV-CB plasmid (a gift from Brian Kaspar) harboring AAV2 ITRs for the production of self-complementary AAV9 (Foust et al., 2010). The corresponding AAV9 vectors packaged into serotype-9 capsid were custom produced by Vector BioLabs using triple plasmid transfection of HEK293 cells and purification by two rounds of CsCl gradient centrifugation. Titer and purity of each AAV9 preparation were confirmed in-house by qPCR (see Table S1 for the sequence of primers) and silver staining, respectively.

For lentiviral production, complementary oligonucleotide templates for an shRNA targeting mouse Stasimon mRNA (5’-GGAAGACCCGTTGTATACA-3’) were annealed and cloned into pSUPERIOR.puro (OligoEngine). To generate the pLenti.pur/Stas_RNAi_ lentiviral vector, a fragment comprising the shRNA cassette under the control of a tetracyline-regulated H1_TO_ promoter and the puromycin resistance gene driven by the PGK promoter were excised from pSUPERIOR.puro and cloned into a modified pRRLSIN.cPPT.PGK-GFP.WPRE vector (Addgene plasmid 12252) lacking the PGK-GFP cassette as a backbone. Viral stocks pseudotyped with the vesicular stomatitis G protein (VSV-G) were prepared by transient co-transfection of HEK293T cells using the ViraPower™ Lentiviral Packaging Mix (Invitrogen) following manufacturer’s instructions.

### Cell lines and treatments

The NIH3T3-Stas_RNAi_ cell line was generated by transduction of NIH3T3 cells with pLenti.pur/Stas_RNAi_, followed by antibiotic selection with 5 μg/ml Puromycin (Sigma) and cloning by limiting dilution as described before (Lotti et al., 2012; Ruggiu et al., 2012).

Mouse NIH3T3 fibroblasts were grown in DMEM with high glucose (Invitrogen) containing 10% of FBS (HyClone), 2 mM glutamine (Gibco), and 10μg/mL Gentamicin (Gibco). NIH3T3 cells were cultured for 4 hours in the presence of 100 nM Bafilomycin A (Alfa Aesar) to block autophagy and/or in DMEM media without glutamine and serum for autophagy induction prior to downstream analysis. For treatment with MOs, each MO (GeneTools) was added to culture media at a final concentration of 10µM along with Endoporter (GeneTools) and incubated for 60 hours. For immunofluorescence analysis following MO treatment, NIH3T3 cells cultured on glass coverslips in 24-well plates were washed briefly with phosphate buffered saline (PBS), fixed with 4% paraformaldehyde (PFA) in PBS for 30 minutes at room temperature, and permeabilized with 0.5% Triton X-100–PBS for 5 minutes on ice. Blocking and incubation with both primary and secondary antibodies were performed using 3% bovine serum albumin (BSA) in PBS for 1 hour at room temperature. Images were collected with an SP5 Leica confocal microscope.

### Animal procedures and behavioral analysis

All surgical procedures on postnatal mice were performed in accordance with the NIH guidelines and approved by the Institutional Laboratory Animal Care and Use Committee of Columbia University. The original breeding pairs for SMNΔ7 (*Smn*^+/-^; *SMN2*^+/+^; *SMNΔ7*^+/+^) mice (stock #005025), PV^Cre^ mice (stock #008069), ChAT^Cre^ mice (stock #006410), and Smn^Res^ (*Smn^Res/+^; SMN2^+/+^; SMNΔ7^+/+^*) mice expressing a Cre-inducible Smn*^Res^* allele (stock #007951) were obtained from Jackson Laboratory. Experimental SMA mice indicated as SMA+SMN (PV^CRE^) and SMA+SMN (ChAT^CRE^) carried one *Smn^Res^* allele and one *Smn* knockout allele, and were heterozygous for either PV^Cre^ or ChAT^Cre^ in the SMNΔ7 background (i.e. *PV^Cre+/−^; Smn^Res/-^; SMN2^+/+^; SMNΔ7^+/+^* and *ChAT^Cre+/−^; Smn^Res/-^; SMN2^+/+^; SMNΔ7^+/+^* respectively). Tail DNA PCR genotyping was performed as described previously (Fletcher et al., 2017). See Table S1 for the sequence of all primers. For AAV9 gene delivery, P0 mice were anesthetized by isoflurane inhalation and injected in the right lateral ventricle of the brain with ∼1x 10^11^ genome copies of AAV9 vectors in a PBS solution containing a vital dye (Fast Green; Sigma-Aldrich). Approximately equal proportions of mice of both sexes were used in all experiments, and aggregated data are presented because gender-specific differences were not found.

For behavioral analysis, mice from all experimental groups were monitored daily, weighed, and three righting reflex tests were timed and averaged as described previously (Mentis et al., 2011). Mice with 25% weight loss and an inability to right were euthanized with carbon dioxide to comply with IACUC guidelines. Righting time was defined as the time for the pup to turn over on its limbs after being placed on its back. The cut-off test time for the righting reflex was 60 seconds to comply with IACUC guidelines.

### Immunohistochemistry

For immunostaining of the spinal cord, mice were sacrificed at P11, the spinal cord was removed natively and post-fixed in 4% PFA overnight at 4°C. On the following day, the spinal cords were briefly washed with PBS, subsequently embedded in warm 5% Agar and serial transverse sections (75 μm) were cut on a Leica VT1000S vibratome. The sections were blocked with 10% normal donkey serum in 0.01 M PBS with 0.3% Triton X-100 (PBS-T; pH 7.4) for 1.5 hours and incubated overnight at room temperature in different combinations of the primary antibodies (see Table S2 for antibody details). The following day, after six 10-minute washes with PBS, secondary antibody incubations were performed for 3 hours with the appropriate species-specific antiserum coupled to FITC, Cy3 or Cy5 (Jackson labs) diluted at 1:250 in PBS-T. After secondary antibody incubations, the sections were washed six times for 10 minutes in PBS and mounted on slides and cover-slipped with an anti-fading solution made of Glycerol:PBS (3:7).

For immunostaining of NMJs, mice were sacrificed at P11 and the QL muscle was dissected and immediately fixed with 4% PFA for 20 minutes. After fixation, single muscle fibers were teased and washed five times in PBS for 10 minutes each followed by staining of the postsynaptic part of the NMJ with α-bungarotoxin coupled to Alexa Fluor 555 in PBS for 20 minutes. Subsequently, the muscle fibers were washed five times in PBS for 10 minutes and blocked with 10% donkey serum in PBS-T for 1 hour. Anti-Neurofilament-M and anti-Synaptophysin 1 antibodies to immunolabel the presynaptic part of the NMJ were applied in blocking solution overnight at 4°C (see Table S2 for antibody details). The muscle fibers were then washed 3 times for 10 minutes in PBS. Secondary antibodies were applied for 1 hour in blocking solution at room temperature. Finally, the muscle fibers were washed 3 times in PBS for 10 minutes and mounted with Vectashield (Vector laboratories, CA) (Simon et al., 2010).

### Confocal microscopy

Confocal microscopy was performed using either SP5 or SP8 (Leica) confocal microscopes as described previously (Simon et al., 2017; Van Alstyne et al., 2018a). Sections were scanned using a 20x or 40x objective. Motor neurons were counted off-line from z-stack images (collected at 3μm intervals in the z axis) from eleven sections of the selected L1 and L5 spinal segments as we previously described (Simon et al., 2017; Van Alstyne et al., 2018a). Only ChAT+ motor neurons that contained the nucleus were counted to avoid double counting from adjoining sections. Quantitative analysis of VGluT1 immunoreactive synaptic densities on motor neurons was performed on image stacks of optical sections scanned using a 40x oil (N.A. 1.25) objective throughout the whole section thickness at 0.4μm z-steps to include the whole cell body and dendrites of ChAT+ motor neurons. The number of VGluT1+ synapses were counted over the entire surface of the motor neuron soma as well as on primary dendrites up to a distance of 100 μm from the soma using Leica LASAF software as previously described (Mentis et al., 2011). For the analysis of muscle innervation, at least 200 randomly selected NMJs per muscle sample were quantified for each biological replicate. Only BTX+ endplates that lacked any pre-synaptic coverage by both synaptophysin and NF-M were scored as fully denervated.

### Electrophysiology

To record the monosynaptic reflex, we conducted the experiment as previously described (Simon et al 2017). The animals were decapitated at P11, the spinal cords dissected and removed under cold (∼12°C) artificial cerebrospinal fluid (aCSF) containing 128.35 mM NaCl, 4 mM KCl, 0.58 mM NaH_2_PO_4_.H_2_0, 21 mM NaHCO_3_, 30 mM D-Glucose, 1.5 mM CaCl_2_.H_2_0, and 1 mM MgSO_4_.7H_2_0. The spinal cord was then transferred to a customized recording chamber. The intact *ex vivo* spinal cord preparation was perfused continuously with oxygenated (95%O_2_/5%CO_2_) aCSF (∼13 ml/min). The dorsal root and ventral root of the L1 segment were placed into suction electrodes for stimulation or recording respectively. The extracellular ventral root potentials were recorded (DC −3kHz, Cyberamp, Molecular Devices) in response to a brief (0.2ms) orthodromic stimulation (A365, current stimulus isolator, WPI, Sarasota, FL) of the L1 dorsal root. The stimulus threshold was defined as the current at which the minimal evoked response was recorded in three out of five trials. Recordings were fed to an A/D interface (Digidata 1440A, Molecular Devices) and acquired with Clampex (v10.2, Molecular Devices) at a sampling rate of 10kHz. Data were analyzed off-line using Clampfit (v10.2, Molecular Devices). The monosynaptic component of the EPSP amplitude was measured from the onset of response to 3ms (Shneider et al., 2009). Measurements were taken from averaged traces of five trials elicited at 0.1Hz. The temperature of the physiological solution ranged between 21-25°C. For synaptic depression experiments, the dorsal root was stimulated at 10Hz for five stimuli and the resulting monosynaptic component of the amplitude recorded and analyzed off-line. The amplitude of the second to the fifth stimulus was expressed as a percentage of the amplitude of the first stimulus.

To functionally assess NMJs of the QL muscle at P11, we applied the same technique as described previously (Fletcher et al., 2017). Motor neurons axons in the ventral root L1 supplying the QL muscle were stimulated by drawing the ventral root into a suction electrode, having removed the spinal cord, and the CMAP was recorded from the muscle using a concentric bipolar electrode. L1 motor neuron axons were stimulated with five stimuli at 0.1Hz. The maximum CMAP amplitude (baseline-to-peak) was measured from five averages.

### RNA and protein analysis

For RNA analysis from NIH3T3 cells and mouse tissue, total RNA purification and RT-qPCR experiments were carried out as previously described (Lotti et al., 2012; Van Alstyne et al., 2018a). The primers used are listed in Table S1. For protein analysis, NIH3T3 cells and mouse tissues were homogenized directly in SDS-PAGE sample buffer and electrophoresed on a 12% SDS-PAGE gel followed by Western blotting as previously described (Ruggiu et al., 2012). The antibodies used are listed in Table S2.

### Statistics

Results are expressed as mean ± standard error of the mean (SEM) from at least three independent experiments using three or more animals per group. Differences between two groups were analyzed by a two-tailed unpaired Student’s *t*-test, whereas differences among three or more groups were analyzed by one-way or two-way ANOVA followed by Tukey’s correction for multiple comparisons as applicable. GraphPad Prism 6 was used for all statistical analyses and P values are indicated as follows: * = P < 0.05; ** = P < 0.01; *** = P < 0.001.

## Acknowledgements

We are grateful to Dr. Brian Kaspar for the kind gift of the dsAAV-CB plasmid. This work was supported by the SMA Foundation (LP and GZM) and NIH grants R01NS078375 (GZM), R01AA027079 (GZM), R21NS077038 (LP) and R01NS102451 (LP).

## Author Contributions

LP designed and supervised the study. CS, MVA., FL, EB and ST performed the experiments and analyzed the data. GZM supervised the electrophysiology studies. CS, MVA and LP wrote the paper with input from all authors.

## Supplemental Figure Legends

**Figure S1.**
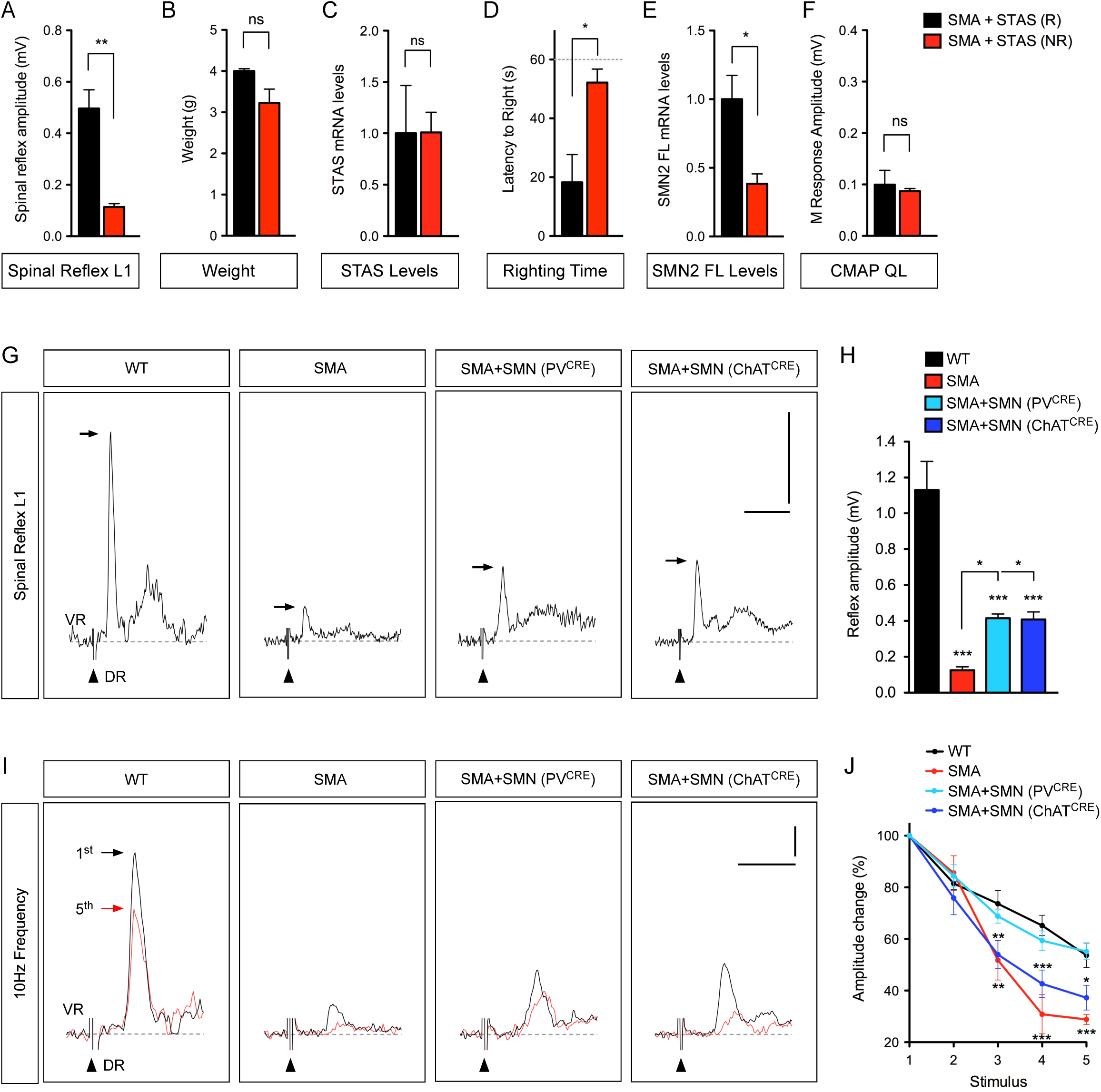
Motor function improvement correlates with enhanced sensory-motor neurotransmission in SMA mice injected with AAV9-STAS. (A) Quantification of spinal reflex amplitudes recorded from SMA mice injected with AAV9-STAS that were either righting (R) or non-righting (NR) at P11. Data represent mean and SEM (n=4). Statistics were performed with two-tailed unpaired Student’s t-test. ** P < 0.01. (B) Body weight in the same groups as in (A) at P11. Data represent mean and SEM (n=4). Statistics were performed with two-tailed unpaired Student’s t-test. ns = not significant. (C) RT-qPCR analysis of human STAS mRNA in spinal cords from the same groups as in (A). Data represent mean and SEM (n=3). Statistics were performed with two-tailed unpaired Student’s t-test. ns = not significant. (D) Righting time in the same groups as in (A) at P11. Data represent mean and SEM (n=4). Statistics were performed with two-tailed unpaired Student’s t-test. * P < 0.05. (E) RT-qPCR analysis of human full-length SMN2 (SMN2 FL) mRNA in spinal cords from the same groups as in (A). Data represent mean and SEM (n=3). Statistics were performed with two-tailed unpaired Student’s t-test. * P < 0.05. (F) M response amplitudes recorded from the QL muscle in the same groups as in (A) at P11. Data represent mean and SEM (n=3). Statistics were performed with two-tailed unpaired Student’s t-test. ns = not significant. (G) Representative traces of extracellular recordings from L1 ventral root (VR) following L1 dorsal root (DR) stimulation from WT and SMA mice as well as SMA mice with selective SMN restoration in either proprioceptive neurons (PV^CRE^) or motor neurons (ChAT^CRE^) at P11. Arrows indicate the maximum amplitude of the monosynaptic reflex. Arrowheads indicate the stimulus artifact. Scale bars = 0.5 mV and 10 ms. (H) Quantification of spinal reflex amplitudes recorded from the same groups as in (G) at P11. Data represent mean and SEM (n≥3). Statistics were performed with one-way ANOVA with Tukey’s *post hoc* test. * P < 0.05; *** P < 0.001. (I) Representative traces of the first (black) and fifth (red) VR responses recorded at P11 following stimulation of the homonymous L1 DR at 10 Hz from the same groups as in (G). Scale bars = 0.2 mV and 5 ms. (J) Quantification of amplitude changes in the monosynaptic reflex following 10 Hz stimulation from the same groups as in (I). Data represent mean and SEM (n≥5). Statistics were performed with two-way ANOVA with Tukey’s *post hoc* test. * P < 0.05; ** P < 0.01; *** P < 0.001.

**Figure S2.**
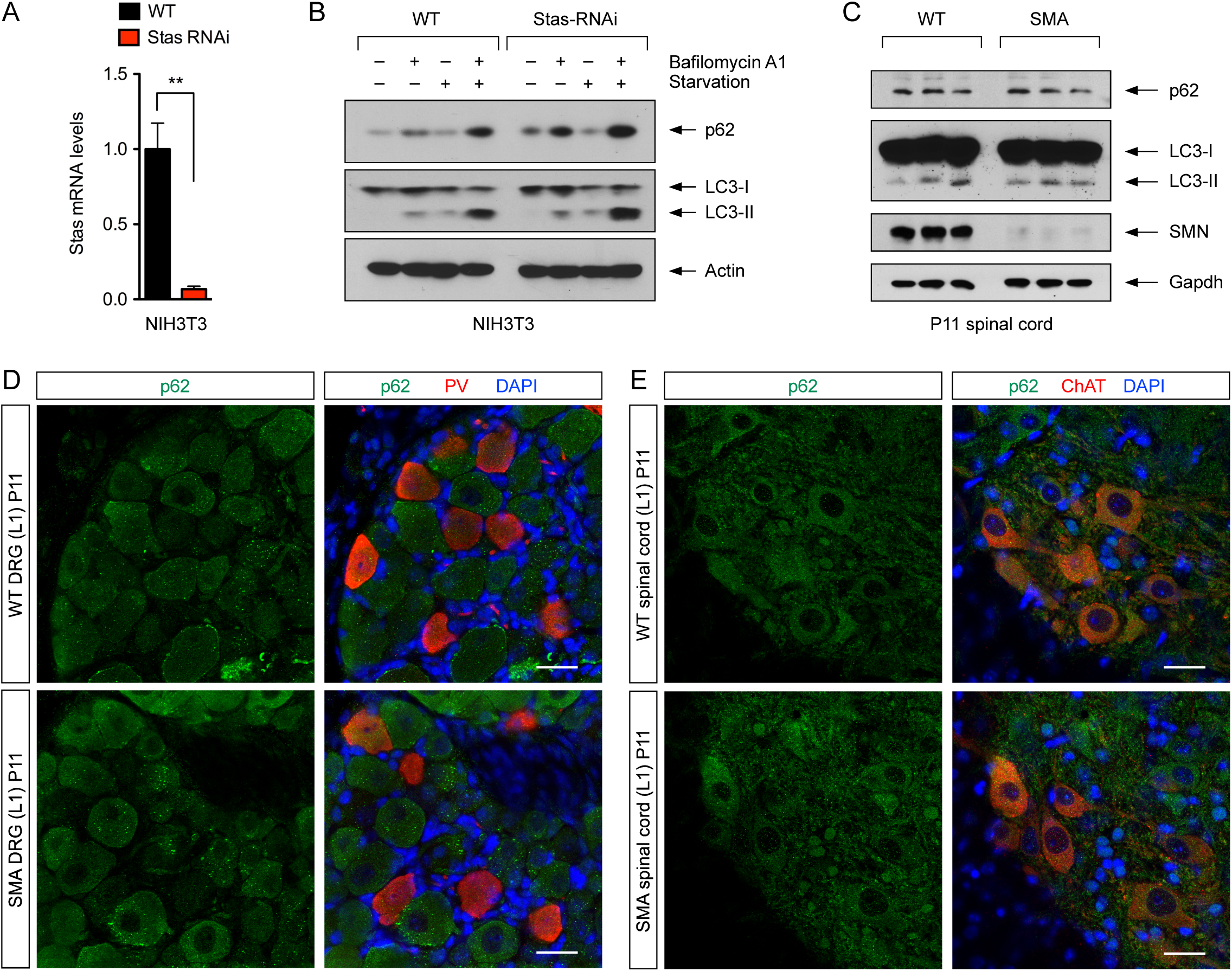
Analysis of autophagy in Stasimon-deficient NIH3T3 cells and SMA mice. (A) RT-qPCR analysis of Stasimon mRNA levels in WT and Stas_RNAi_ NIH3T3 cells. Data represent mean and SEM (n=3). Statistics were performed with two-tailed unpaired Student’s t-test. ** P < 0.01. (B) Western blot analysis of the autophagy markers p62 and LC3 in WT and Stas_RNAi_ NIH3T3 cells that were either untreated or treated with Bafilomycin A1 and serum starvation as indicated. (C) Western blot analysis of p62 and LC3 in spinal cords of WT and SMA mice at P11. Three biological replicates per groups are shown. (D) Immunostaining of L1 DRGs from WT and SMA mice at P11 with p62 (green) and PV (red). Nuclei were counterstained with DAPI (blue). Scale bar= 25 µm. (E) Immunostaining of L1 spinal cords from WT and SMA mice at P11 with p62 (green) and ChAT (red). Nuclei were counterstained with DAPI (blue). Scale bar= 25 µm.

**Table S1.**
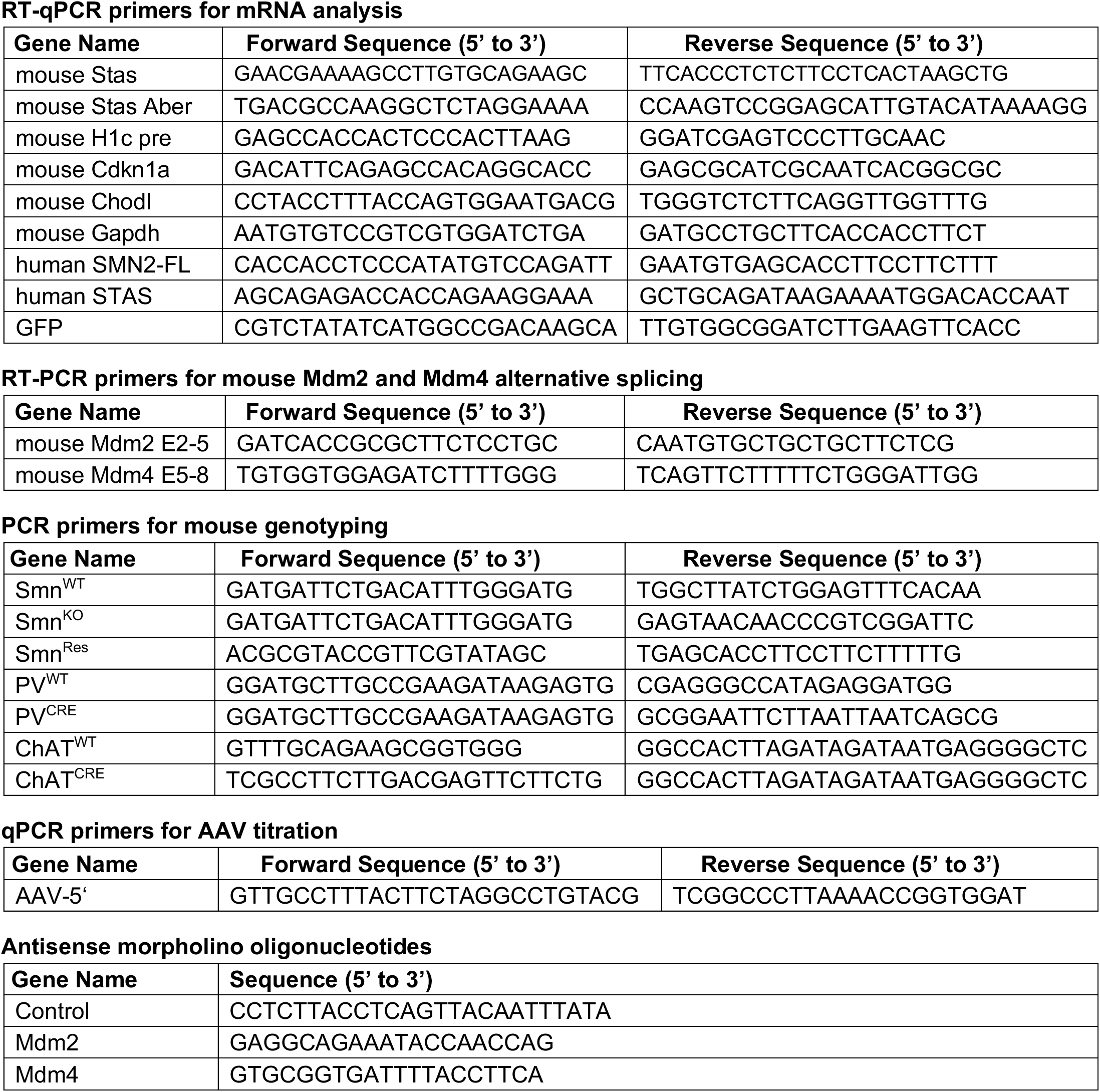
List of primers used in this study.

**Table S2.** List of antibodies used in this study.

| Name | Source | Cat # | Host | Application | Dilution |
| --- | --- | --- | --- | --- | --- |
| SMN (clone 8) | BD Transd Lab | 610646 | Mouse | WB | 1:10,000 |
| GFP | Sigma | G1544 | Rabbit | WB | 1:5,000 |
| p62 | MBL | PM045 | Rabbit | WB | 1:1,000 |
| LC3B | Novus | NB600-1384 | Rabbit | WB | 1:4,000 |
| Actin | Proteintech | 66009 | Mouse | WB | 1:10,000 |
| Gapdh | Millipore | MAB374 | Mouse | WB | 1:5,000 |
| Tubulin (DM1A) | Sigma | T9026 | Mouse | WB | 1:10,000 |
| p62 | MBL | PM045 | Rabbit | IF | 1:500 |
| p53 | Leica Novocastra | NCL-p53-CM5p | Rabbit | IF | 1:1,000 |
| p-p53<sup>S15</sup> | Cell Signaling | 9284 (Lots: 12,15) | Rabbit | IF | 1:200 |
| Parvalbumin | Covance | custom made | Chicken | IF | 1:10,000 |
| VGluT1 | Covance | custom made | Guinea pig | IF | 1:5,000 |
| Synaptophysin | Synaptic Systems | 101-004 | Guinea pig | IF | 1:500 |
| NF-M | Millipore | AB1987 | Rabbit | IF | 1:1000 |
| ChAT | Millipore | AB144P | Goat | IF | 1:100 |
| Bungarotoxin | Invitrogen | T1175 | N/A | IF | 1:500 |
| DAPI | Molecular Probes | D3571 | N/A | IF | 1:1,000 |
| 488 Anti-rabbit | Jackson | 711-545-152 | Donkey | IF | 1:250 |
| 488 Anti-goat | Jackson | 705-545-147 | Donkey | IF | 1:250 |
| Cy3 Anti-rabbit | Jackson | 711-165-152 | Donkey | IF | 1:250 |
| Cy3 Anti-mouse | Jackson | 715-165-150 | Donkey | IF | 1:250 |
| Cy3 Anti-goat | Jackson | 705-165-147 | Donkey | IF | 1:250 |
| Cy5 Anti-goat | Jackson | 705-175-147 | Donkey | IF | 1:250 |
| Cy5 Anti-guinea pig | Jackson | 706-175-148 | Donkey | IF | 1:250 |
| HRP Anti-mouse | Jackson | 115-035-044 | Goat | WB | 1:10,000 |
| HRP Anti-rabbit | Jackson | 111-035-003 | Goat | WB | 1:5,000 |

